# Optogenetic inhibition of the caudal substantia nigra inflates behavioral responding to uncertain threat and safety

**DOI:** 10.1101/2023.02.18.529041

**Authors:** Kristina M Wright, Shannon Cieslewski, Amanda Chu, Michael A McDannald

**Author notes:** Correspondence: Dr. Michael A. McDannald, 140 Commonwealth Avenue, Chestnut Hill, MA 02467.

## Abstract

Defensive responding is adaptive when it approximates current threat, but maladaptive when it exceeds current threat. Here we asked if the substantia nigra, a region consistently implicated in reward, is necessary to show appropriate levels of defensive responding in Pavlovian fear discrimination. Rats received bilateral transduction of the caudal substantia nigra with halorhodopsin or a control fluorophore, and bilateral ferrule implants. Rats then behaviorally discriminated cues predicting unique foot shock probabilities (danger, *p*=1; uncertainty, *p*=0.25; and safety, *p*=0). Green-light illumination (532 nm) during cue presentation inflated defensive responding of halorhodopsin rats – measured by suppression of reward seeking – to uncertainty and safety beyond control levels. Green-light illumination outside of cue presentation had no impact on halorhodopsin or control rat responding. The results reveal caudal substantia nigra cue activity is necessary to inhibit defensive responding to non-threatening and uncertain threat cues.

## Introduction

The display of defensive behavior is adaptive in the face of potential threat (Bouton & Bolles, 1980; Estes & Skinner, 1941). Rather than being absolute, the degree to which defensive behavior is elicited can scale to degree of threat (Ray et al., 2020; Rescorla, 1968). Individuals with stress and anxiety disorders typically acquire defensive responding to threat cues, but can have difficulty limiting responding to threat (Christianson et al., 2012; Jovanovic et al., 2012). Uncovering neural circuits that permit defensive responding to approximate degree of threat may provide insight into healthy and disordered defensive responding.

Long studied in rewarding contexts (Han et al., 1997; Schultz et al., 1993), there is increasing evidence of a role for the substantia nigra in Pavlovian fear conditioning (Baldi et al., 2007; Kinoshita et al., 2015). Notably, Bouchet et al. (2018) found that substantia nigra dopamine neurons function to reduce defensive responding (freezing) in an extinction setting. Rats received Pavlovian fear conditioning, then excitatory, chemogenetic stimulation of substantia nigra neurons during extinction. Rats receiving chemogenetic stimulation showed enhanced recall of extinction (reduced freezing) and no renewal of responding (no freezing) (Bouchet et al., 2018). These findings mark the substantia nigra as a key brain region to reduce defensive responding.

The findings raise further questions. Most pertinent, is substantia nigra activity *during* cue presentation necessary to show appropriate defensive responding? To answer, we took a within-subject’s, optogenetic inhibition approach (Ray et al., 2020). Male, Long Evans rats received bilateral transduction of the caudal substantia nigra with halorhodopsin (eNpHR) or a control fluorophore (YFP). Rats received Pavlovian fear discrimination in which three cues predicted unique foot shock probabilities: danger (*p*=1), uncertainty (*p*=0.25), and safety (*p*=0). After discrimination was established, eNpHR and YFP rats received green-light illumination during cue presentation or during inter-trial intervals. If substantia nigra cue activity is necessary to show appropriate defensive responding to cues, then only eNpHR rats receiving cue illumination should increase responding.

## Materials and Methods

### Subjects

Subjects were 17 male Long Evans rats approximately 60 days old on arrival, obtained from Charles River Laboratories, and maintained on a 12-hr light cycle (lights off at 6:00 PM). Rats were individually housed and acclimated to the animal facility with food and water freely available for three days. Following acclimation, rats were restricted to and maintained at 85% of their free-feeding body weight. All rats were returned to ad *libitum food,* received surgery, recovered, and were again maintained at 85% of their free-feeding body weight for the duration of behavioral testing. All protocols were approved by the Boston College Animal Care and Use Committee and all experiments were carried out in accordance with the NIH guidelines regarding the care and use of rats for experimental procedures.

### Surgical procedures

Aseptic, stereotaxic surgery was performed under isoflurane anesthesia (1–5% in oxygen). Rimadyl (subcutaneous, 5 mg/kg), lidocaine (subcutaneous, 2%), and lactated ringer’s solution (~2–5 mL) were administered preoperatively. The skull was exposed via midline incision and scoured in a crosshatch pattern with a scalpel blade to increase resin adhesion. Nine holes were drilled: five for screws, two for infusion and two for ferrules. Five screws were installed in the skull to stabilize the connection between the skull, bilateral optical ferrule implants and a protective head cap (screw placements: two anterior to bregma, two between bregma and lambda about ~3 mm medial to the lateral ridges of the skull, and one on the midline ~5 mm posterior of lambda). Infusions were delivered at a rate of ~0.11 μl/min, using a 2 μl Neuros syringe controlled by a microsyringe pump. Rats received bilateral 0.5 μl infusions of halorhodopsin, AAV5-hSyn-eNpHR3.0-YFP (n = 9) or a control fluorophore AAV5-hSyn-EYFP (n = 8) aimed at the caudal substantia nigra (caudal substantia nigra): AP −7.10mm, ML +/- 1.90mm, DV −7.75mm. Bilateral optical ferrules were implanted dorsal to the caudal substantia nigra at a 15° angle: AP −6.85mm, ML +/- 3.08mm, DV −6.50mm. Ferrule implants were protected by a black, light-occluding head cap made from a modified 50mL falcon tube. The head cap and ferrules were cemented to the skull using orthodontic resin. Post-surgery, rats received 8-12 days of undisturbed recovery and 14 days of oral Cephalexin mixed with Froot Loops to encourage consumption. Dust caps protected the ends of optical ferrule implants during recovery and all behavior sessions when fiber optic cables were not in use.

### Optogenetic ferrule and fiber optic cable assembly

Optical ferrules were constructed using 2.5mm Ceramic Dome Ferrule Assemblies: 230um ID, bore tolerance: −0/+10um, Concentricity < 20um paired with multimode optical fiber, 0.22 NA, High-OH, Ø200 μm Core for 250 - 1200 nm. Ferrules were assembled, polished, and inspected for flares. Light output was tested with a Si Sensor Power Meter and a 532nm, 499 mW green laser identical to those used for light illumination during behavior testing. Ferrules were polished until 30-40mW of light from the laser source could produce at least 25mW of light output from an attached ferrule. Source laser mW requirements (to achieve 25mW ferrule output) were matched for each ferrule pair implanted during surgery. This way, the same amount of source laser light would result in equivalent light intensities in each hemisphere. Bilateral behavior cables consisted of a single metal-shielded shaft encompassing two cladded multimode optical fibers, 0.39 NA, High-OH, Ø200 μm Core for 300 - 1200 nm, TECS Clad. A wye splitter at each end separated each fiber from the central shaft to accommodate individual ferrule-to-ferrule connections with implants on the rat’s head and multimode FC connections to the 1×2 rotary commutator above the experimental chamber. Following fabrication, all cables were re-tested prior to each illumination session to ensure that there was a difference of no more than 5 mW of laser output between each side. Finally, the source laser power was calibrated for each light illumination session, so that the final cable output would pass the amount of light required for paired ferrule implants to permit either 12.5 mW or 25 mW light delivery into the brain.

### Behavioral apparatus

The apparatus consisted of four individual experimental chambers (internal dimensions: 30.5 cm × 24.1 cm × 29.2 cm) with aluminum front and back walls, clear acrylic sides and top, and a grid floor (0.48 cm diameter bars spaced 1.6 cm apart). Each grid floor bar was electrically connected to an aversive foot shock generator. An external food cup was present at the center of one wall 2.5 cm above the grid floor. A central panel nose poke opening, equipped with infrared photocells (sampled at approximately 1 kHz), was centered 8.5 cm above the food cup (apparatus visually summarized in Figure 1A). Each experimental chamber was enclosed in a sound-attenuating shell.

**Figure 1.**
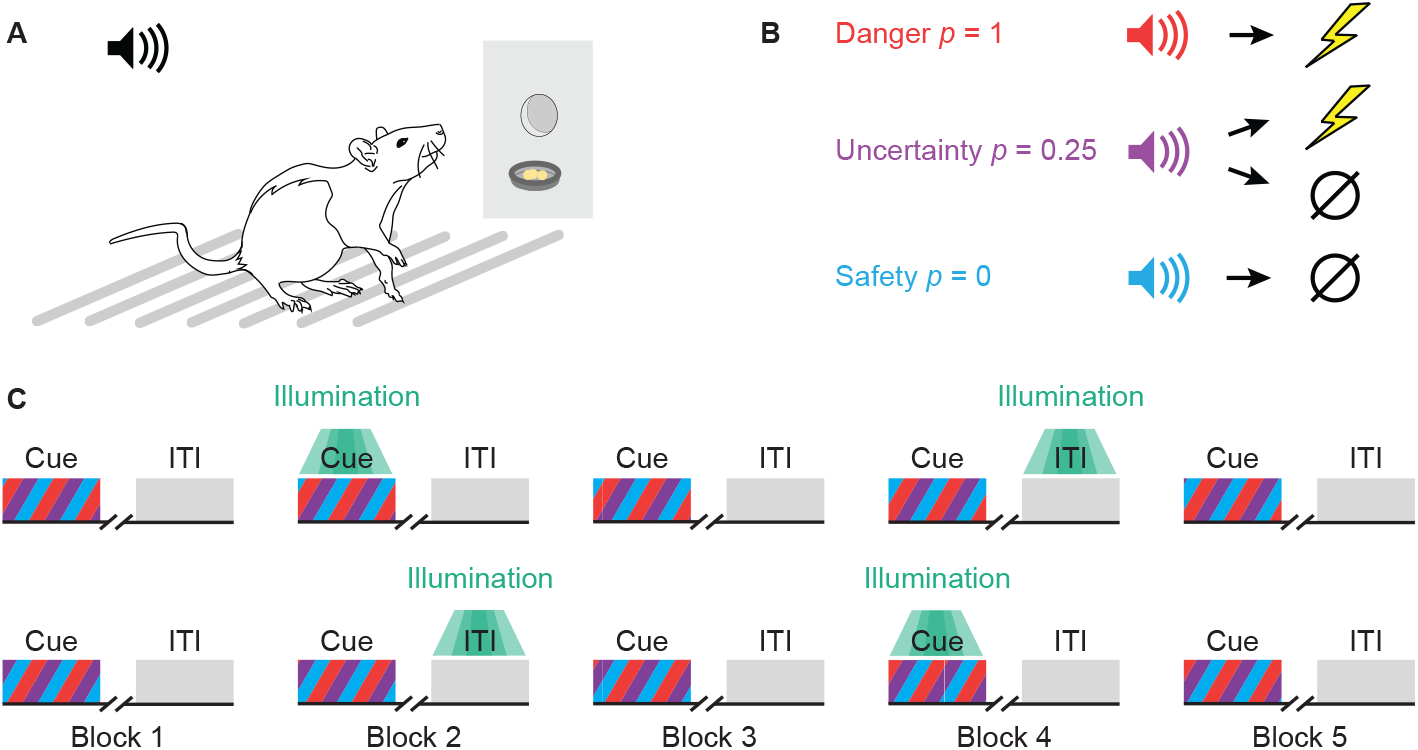
Experimental outline. (**A**) Behavioral testing took place in an experimental chamber equipped with overhead speaker, grid floor, central nose poke port, and an external food cup. (**B**) Pavlovian fear discrimination consisted of three auditory cues predicting unique foot shock probabilities: danger, p = 1 (red); uncertainty, p = 0.25 (purple); and safety (cyan), p = 0. (**C**) Green-light illumination (532 nm) was administered for 10 s during cue presentation (Cue Illumination) or during the inter-trial interval (ITI Illumination). All rats receiving Cue and ITI illumination over a 5-block sequence. Illumination order was counterbalanced, with half of the rats receiving Cue first (top) and half ITI first (bottom).

Green lasers (532nm) were used for illumination. A 5-inch diameter hole in the chamber ceiling funneled to a ~1.5 inch whole just below the commutator, permitting fiber optic cables to be threaded into the experimental chamber from above, and allowed them to move freely with each animal during optogenetic behavior sessions. Fiber optic cables were suspended from a 1 × 2 fiber optic rotary commutator mounted to the shell ceiling. Two speakers were mounted 20 cm apart on the shell ceiling. Chambers were illuminated with a small strip of red LED lights mounted on the shell ceiling.

### Behavioral procedures

#### Pellet exposure

Each rat was exposed to 4 grams of reward pellets in their home cage on two days, followed by one day of automatic pellet delivery to the food cup inside the experimental chamber.

#### Nose poke acquisition

Each rat was shaped to nose poke for pellet delivery using a fixed ratio schedule in which one nose poke yielded one pellet. Nose poke acquisition sessions lasted for 30 minutes or until approximately 50 nose pokes were completed. Rats moved on to variable interval (VI) schedules in which nose pokes were reinforced on average every 30 s (day 1), or 60 s (days 2-5). For the remainder of behavioral testing, nose pokes were reinforced on a VI-60 schedule independent of all Pavlovian contingencies.

#### Cue pre-exposure

Each rat was pre-exposed to the three auditory cues to be used in Pavlovian discrimination in two, 42-minute sessions. The 10 s auditory cues were repeating, 500 ms motifs of a horn, siren or broadband click. Previous studies have found these cues to be equally salient, yet readily discriminable (Chu et al., 2022; Strickland & McDannald, 2022; Wright & McDannald, 2019). Sessions consisted of four presentations of each cue (12 total presentations) with a mean inter-trial interval of 3.5 min. Trial type order was randomly determined by the behavioral program, and differed for each rat during each session.

#### Pavlovian fear discrimination

Following pre-exposure, all rats received 8, 64-minute behavior-only discrimination sessions. A single session consisted of 18 cue trials: four danger trials, six uncertainty no-shock trials, two uncertainty shock trials, and six safety trials, with a mean inter-trial interval of 3.5 min. Each auditory cue was associated with a unique foot shock probability (0.5 mA, 0.5 s): danger, *p* = 1; uncertainty, *p* = 0.25; and safety, *p* = 0 (Figure 1B). The physical identities of the cues were counterbalanced. Foot shock was administered two seconds following the termination of the auditory cue.

#### Green-light illumination

The remaining 10 discrimination sessions were divided into 5, 2-session blocks. All rats were habituated to optogenetic cables during block 1. For Cue-ITI rats [(eNpHR (n = 3), YFP (n = 4)] blocks 2-5 were Cue illumination, no illumination, ITI illumination and no illumination. For ITI-Cue rats [(eNpHR (n=6), YFP (n=4)] blocks 2-5 were ITI illumination, no illumination, Cue illumination and no illumination (Figure 1C).

During Cue illumination sessions, 12.5 mW or 25 mW, 532 nm green light was delivered bilaterally for the entirety of all 10 s cues. Two light levels were chosen to observe possible dose-dependent effects of intensity. For example, 12.5 mW may be insufficient to inhibit activity to alter behavior but 25 mW may be sufficient. During ITI illumination sessions, light was delivered for 10 s ITI periods between cue trials. No illumination sessions provided measures of pre and post illumination responding for comparison to illumination trials. Rats were not plugged into behavior cables during no-illumination blocks 3 and 5. Due to a programming error, ITI illumination sessions contained one additional 10-second illumination (versus Cue illumination sessions) for a total of 19, 10s illumination periods.

#### Histology

Rats were deeply anesthetized using isoflurane, perfused with 0.9% biological saline and 4% paraformaldehyde in a 0.2 M potassium phosphate buffered solution. Brains were extracted, post-fixed in 10% neutral-buffered formalin for 24 hr, stored in 10% sucrose/formalin and sectioned via microtome. All brains were processed for fluorescent microscopy using anti-tyrosine hydroxylase immunohistochemistry (Tyrosine hydroxylase primary paired with Alexa 594 secondary) and NeuroTrace.

#### Statistical analyses

Behavioral data were acquired using Med Associates, Med-PC IV software. Raw data were processed in Matlab to extract time stamps for nose pokes, cues, foot shocks and illumination. Baseline nose poke rate was the mean of the 20 s prior to cue presentation. Cue nose poke rate was the mean of the 10-s cue. Suppression of rewarded nose poking was calculated as a ratio: (baseline poke rate - cue poke rate) / (baseline poke rate + cue poke rate). A ratio of 1 indicated complete nose poke suppression during cue presentation, while 0 indicated continued pressing at baseline rates. Gradations between 1 and 0 indicated intermediate levels of cue-elicited nose poke suppression.

Suppression ratios were analyzed using repeated measures ANOVA in SPSS with between factors of group (eNpHR/YFP), intensity (12.5 mW or 25 mW of laser illumination) and order (Cue versus ITI illumination first); within factors of cue (danger, uncertainty, safety), block (2-session blocks 1 through 5) & illumination (Cue versus ITI laser illumination). Post-hoc tests were performed using 95% bootstrap confidence intervals (BCIs). For ANOVA analyses, p < 0.05 was considered significant. 95% BCIs were preferred over t-tests because they do not assume normality and avoid issue with multiple comparisons. For 95% BCIs, differences were reported if the confidence interval did not contain zero.

## Results

### Histology

The caudal substantia nigra was successfully transducted in YFP and eNpHR rats (Figure 2A). While somewhat diffuse, transduction was concentrated in tyrosine hydroxylase-containing regions substantia nigra compacta (dorsal tier) and reticulata, with each individual showing greater than 90% YFP expression in the caudal substantia nigra at Bregma −6.60 mm. Areas of most consistent transduction for each group of rats are the deepest yellow: between Bregma −6.12 mm and −6.84 mm for YFP, or Bregma −5.88 mm and −6.60 mm for eNpHR rats. Ferrule placements were confirmed to be dorsal to the caudal substantia nigra, at Bregma −6.36 mm +/- 0.72 mm. Each rat’s complete transduction and accompanying ferrule tip placements were drawn from fluorescent slices processed for tyrosine hydroxylase immunohistochemistry, made translucent and stacked (Figure 2B).

**Figure 2.**
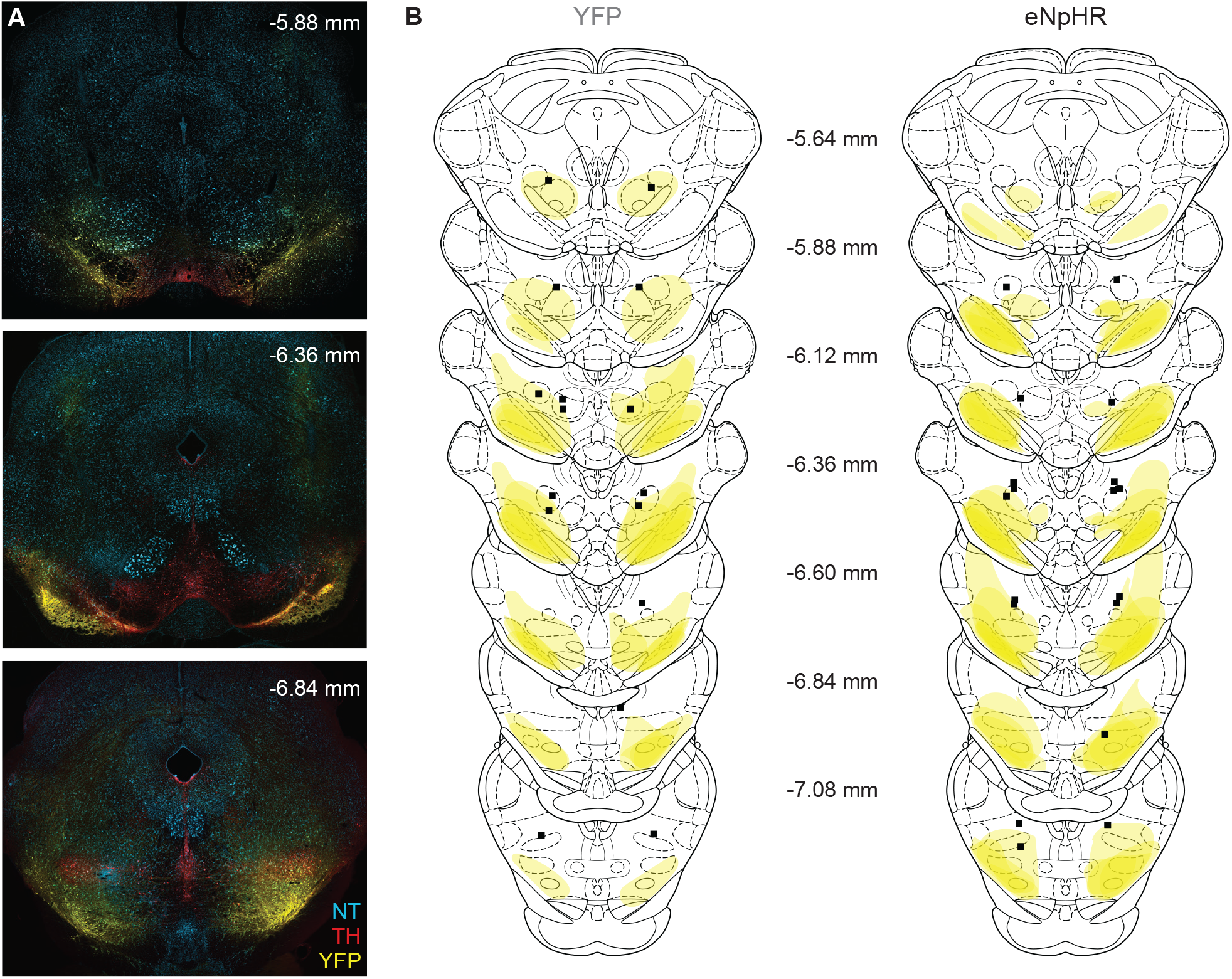
Histology. (**A**) Representative expression of YFP (yellow) tyrosine hydroxylase (red), and NeuroTrace (blue) is shown for 3 bregma levels. (**B**) Extent of transduction and fiber optic ferrule placements are plotted for all rats YFP (n = 8, left) and eNpHR (n = 9, right) across 7 bregma levels.

### Baseline nose poke rates

YFP and eNpHR rats had equivalent baseline nose poke rates throughout testing (Figure 3A). ANOVA for baseline nose poke rate [between factors: group (YFP vs. eNpHR), intensity (12.5 mW vs. 25 mW) and order (ITI-Cue vs. Cue-ITI); within factors: session (18)] found a significant main effect of session (F_17,170_ = 3.03, *p* = 1.27 × 10^-4^), as well as a trend toward significance for a group x session interaction (F_17,170_ = 1.66, *p* = 0.056). However, ANOVA for baseline nose poke rate excluding intensity and order found only a main effect of session (F_17,255_ = 3.03, *p* = 9.00 × 10^-6^). No main effect of group was detected in either ANOVA. These results minimize concerns that group differences in cue-elicited, nose poke suppression are the result of differences in baseline rewarded nose poking.

**Figure 3.**
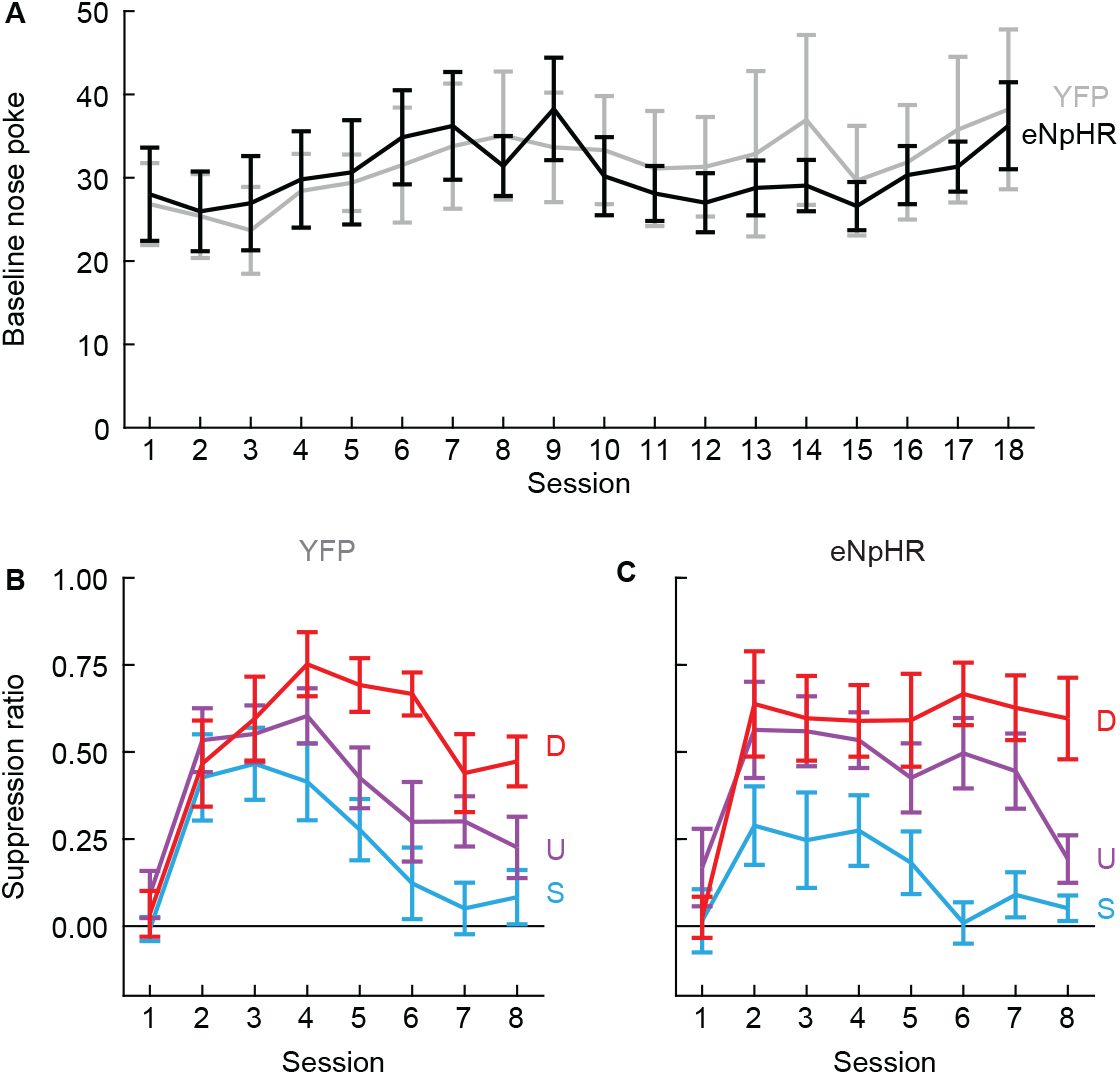
Pre-illumination responding. (**A**) Mean ± SEM baseline nose poke rate is shown YFP rats (gray) and eNpHR rats (black) for the 18 sessions of behavioral testing. (**B**) Mean ± SEM suppression ratio for the 8, pre-illumination discrimination sessions is shown for YFP rats for danger (red), uncertainty (purple), and safety (cyan). (**C**) Mean ± SEM suppression ratio for the 8, pre-illumination discriminations sessions is shown for eNpHR rats, colors as in B.

### Initial fear discrimination

Behavioral discrimination was observed by the eighth session. Suppression ratios were high to danger, intermediate to uncertainty, and low to safety (Figure 3B, C). YFP (Figure 3B) and eNpHR rats (Figure 3C) acquired equivalent discrimination prior to light illumination. ANOVA for suppression ratios [between factor: group (YFP vs. eNpHR); within factors: cue (danger vs. uncertainty vs. safety) & session (8)] found a significant effect of cue (F_2,30_ = 37.12, *p* = 7.73 × 10^-9^), session (F_7,105_ = 18.07, *p* = 1.41 × 10^-15^), and a cue x session interaction (F_14,210_ = 5.47, *p* = 6.48 × 10^-9^). ANOVA revealed no significant main effect of or interaction with group (Fs < 1.6, *p*s > 0.2). Differences in behavioral responding during illumination cannot be attributed to pre-existing differences prior to illumination.

### Optogenetic inhibition during cue presentation inflates responding to uncertainty and safety

A causal role for the caudal substantia nigra requires that changes in cue responding are specific to eNpHR rats receiving Cue illumination. We organized suppression ratio data for danger, uncertainty, and safety into 3, 2-session blocks. Block 1 (pre-illumination) served as a discrimination baseline, no green-light illumination occurred during these sessions. Block 2 (illumination) showed cue responding during the illumination sessions. For Cue sessions green-light illumination occurred during cue presentation (Figure 4A, B). For ITI sessions green-light illumination occurred during the inter-trial interval, at least 1 min after cue presentation (Figure 4C, D). Block 3 (post-illumination) again contained no green-light illumination. The within-subject design meant that individual, cue responding during Cue and ITI illumination could be directly compared.

**Figure 4.**
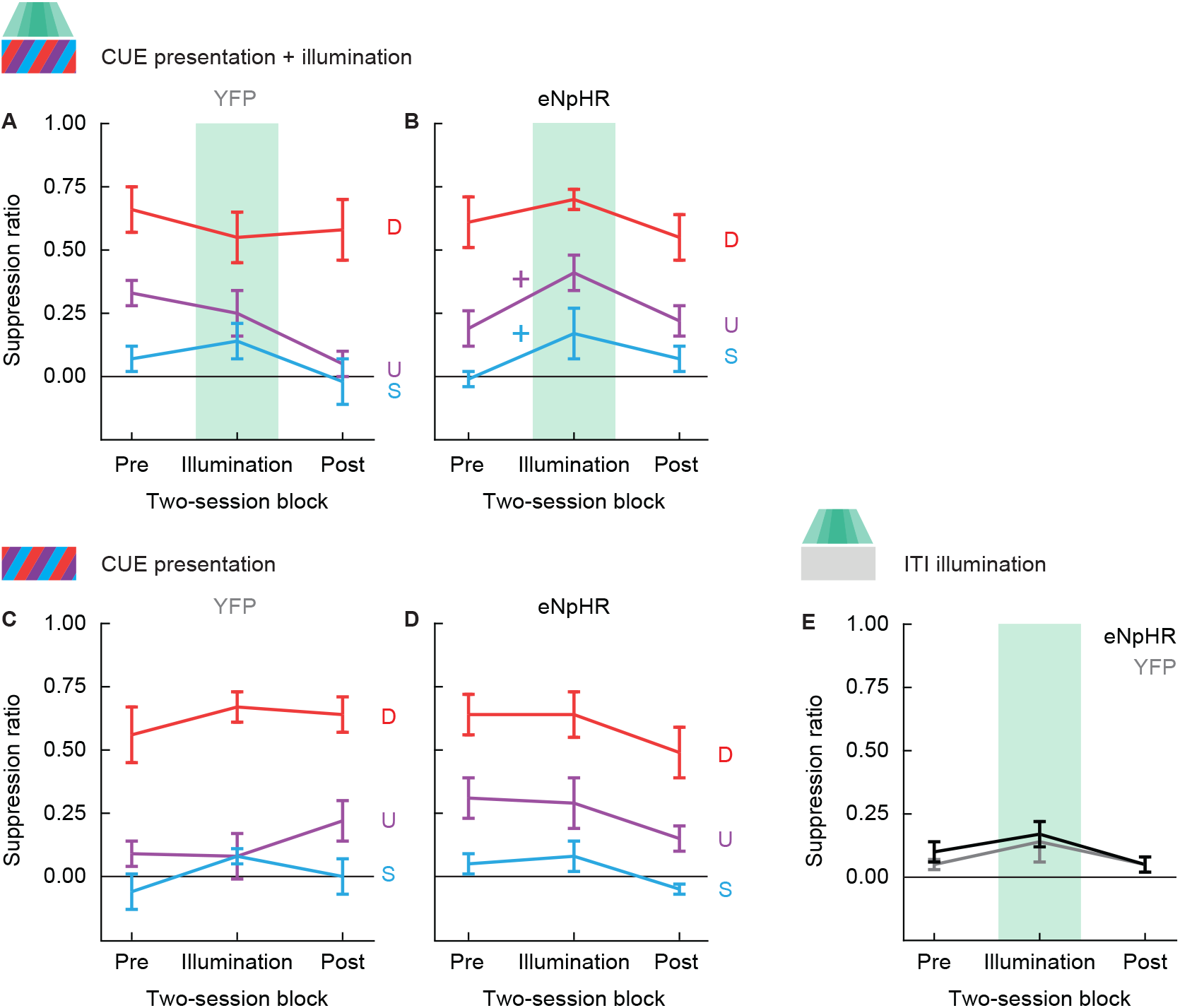
Illumination responding. (**A**) Mean ± SEM suppression ratio for pre-illumination, Cue illumination and post-illumination discrimination sessions is shown for YFP rats for danger (red), uncertainty (purple), and safety (cyan). Green box indicates the block during which Cue green-light illumination occurred. (**B**) Mean ± SEM suppression ratio for pre-illumination, Cue illumination and post-illumination discrimination sessions is shown for eNpHR rats, colors as in A. (**C**) Mean ± SEM suppression ratio for pre-illumination, ITI illumination and post-illumination discrimination sessions is shown for YFP rats for danger (red), uncertainty (purple), and safety (cyan). (**D**) Mean ± SEM suppression ratio for pre-illumination, ITI illumination and post-illumination discrimination sessions is shown for eNpHR rats, colors as in C. (**E**) Mean ± SEM suppression ratio for pre-illumination, ITI illumination and post-illumination discrimination sessions is shown for the inter-trial interval, green-light illumination period for YFP rats (gray) and eNpHR rats (black). Green box indicates the block during which ITI green-light illumination occurred. +95% bootstrap confidence interval for (illumination - pre-illumination) difference score does not contain zero. Color indicates cue: uncertainty (purple) and safety (cyan).

We performed ANOVA for suppression ratio with all factors [between factors: group (YFP vs. eNpHR), intensity (12.5 mW vs. 25 mW) and order (ITI-Cue vs. Cue-ITI); within factors: illumination (Cue vs. ITI), cue (danger vs. uncertainty vs. safety) & block (3, 2-session blocks: pre vs. illumination vs. post)]. Complete ANOVA results are reported in Table 1. Intensity and order were not major determinants of responding, with each involved in only 3 significant interactions. To avoid spurious significance we removed intensity and order, performing ANOVA with factors of group, illumination, cue, and block. ANOVA revealed a significant main effect of cue (F_2,30_ = 96.67, *p* = 8.37 × 10^-14^), indicating discrimination across all conditions. Of most interest, ANOVA revealed a significant group x illumination x block interaction (F_2,30_ = 6.79, *p* = 0.004). The interaction was driven by increased suppression ratios during the illumination block in eNpHR rats receiving Cue illumination (Figure 4B). To identify cue-specific changes in suppression ratios we constructed 95% bootstrap confidence intervals (BCIs) for difference scores (illumination suppression ratio – pre-illumination suppression ratio) for each cue. Observing 95% BCIs that do not contain zero support changes in cue-elicited suppression from the pre-illumination block to the illumination block. 95% BCIs for eNpHR rats receiving Cue illumination revealed increased suppression ratios to uncertainty (mean = 0.22, 95% BCI [(lower bound) 0.09, (upper bound) 0.35]), and safety (M = 0.18, 95% CI [0.004, 0.35]). Changes in suppression ratio to danger were not observed, most likely due to a ceiling effect. 95% BCIs contained zero for all other cues in all other conditions (eNpHR-ITI, YFP-Cue, and YFP-ITI), indicating no change from pre-illumination to illumination sessions.

**Table 1.**
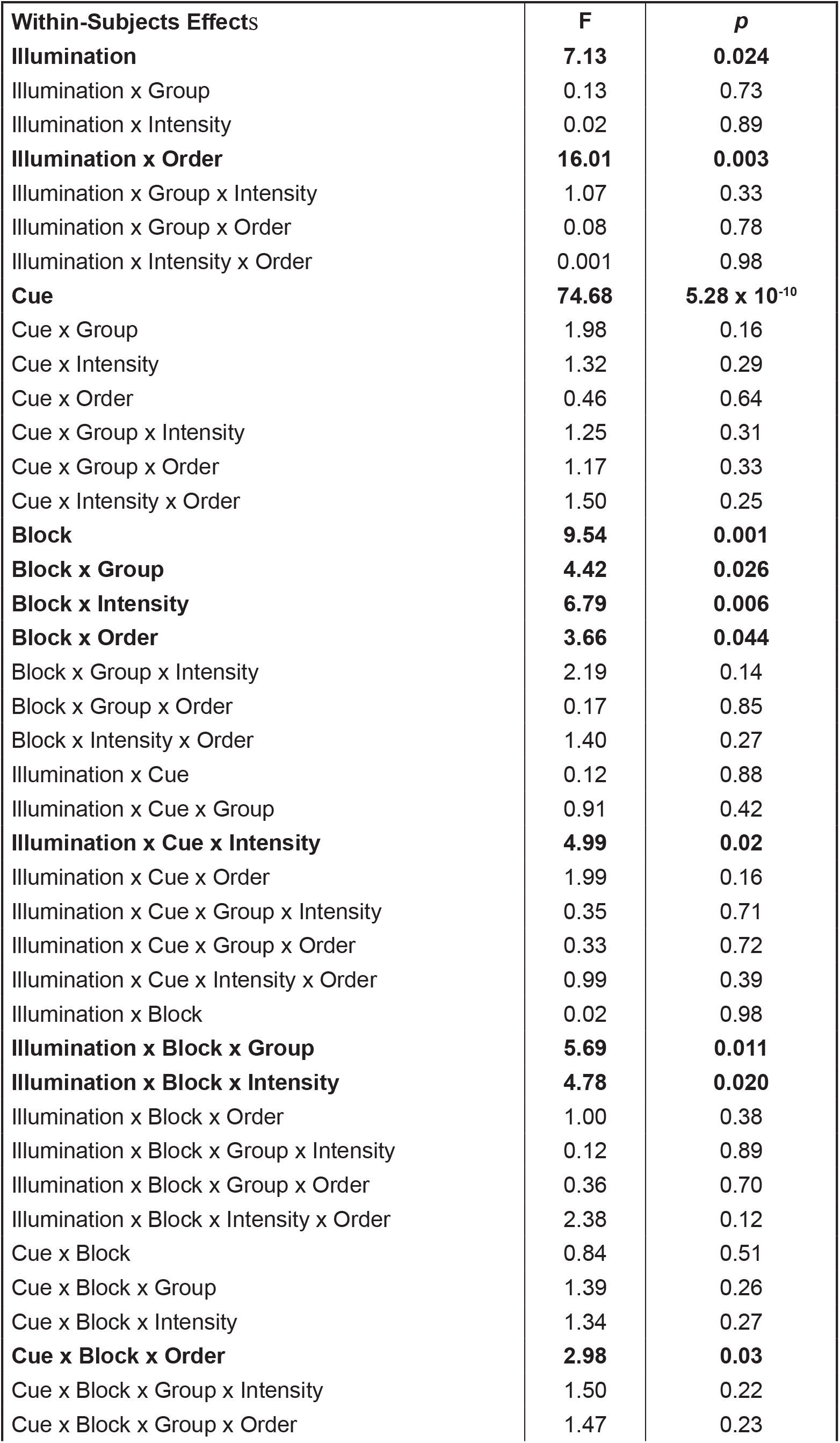
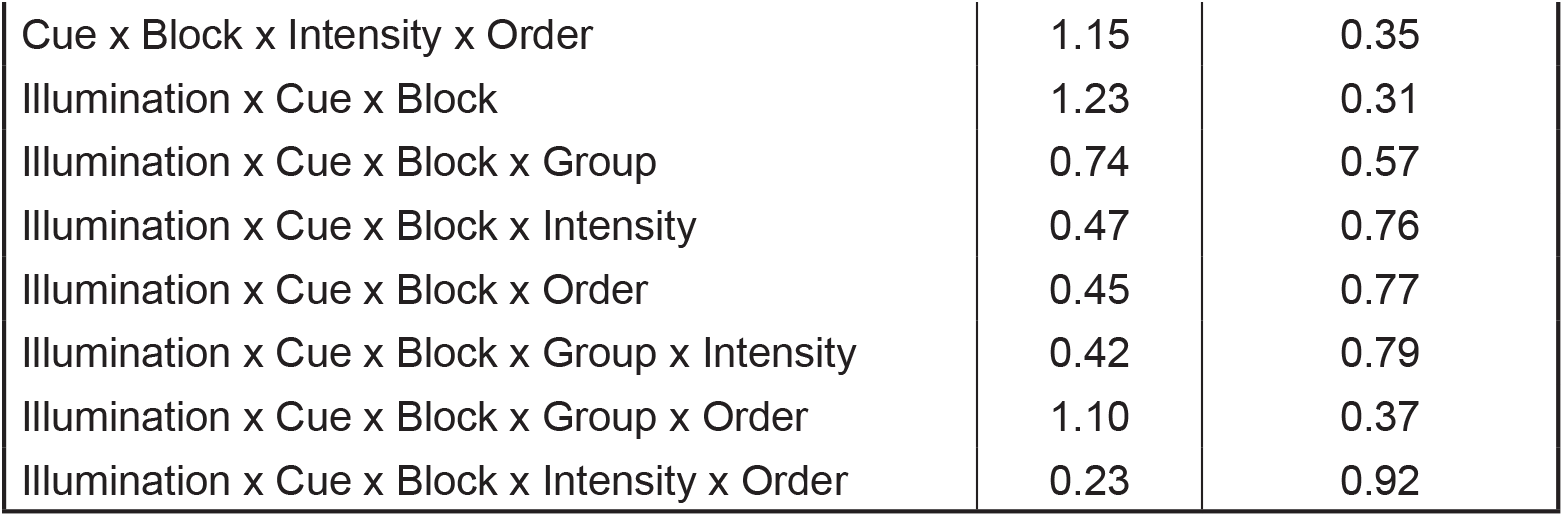
Complete ANOVA results for illumination sessions.

### Illumination-induced inflation of responding is specific to cue presentation

ANOVA and 95% BCI results support the interpretation that optogenetic inhibition of the caudal substantia nigra inflates cue-elicited suppression of reward seeking. However, it is possible that optogenetic inhibition in the absence of cue presentation would be sufficient to suppress reward seeking. To determine this, we examined behavioral responding during the inter-trial interval over the 3-block, ITI illumination sequence (Figure 4E). Suppression ratio data were organized into pre illumination, ITI illumination, and post illumination blocks. Suppression ratios were calculated for the 10-s green-light illumination period during the ITI block, and for comparable, empty periods during the pre-illumination and post-illumination blocks. ANOVA for suppression ratio with all factors [between factors: group (YFP vs. eNpHR), intensity (12.5 mW vs. 25 mW) and order (ITI-Cue vs. Cue-ITI); within factors: block (3, 2-session blocks: pre vs. illumination vs. post)] found no significant main effects or interactions (all F < 1.6, all *p* > 0.2). Neither green-light illumination nor optogenetic inhibition alone was insufficient to suppress reward seeking.

## Discussion

We found that inhibiting caudal substantia nigra activity during cue presentation in a Pavlovian fear discrimination setting increased cues ability to suppress reward seeking. Responding increases were most apparent to the uncertainty and safety cues, and were not observed when activity was inhibited outside of cue presentation. The results support and extend those of Bouchet et al. (2018), further revealing the substantia nigra as a key suppressor of defensive responding.

Two limitations must be pointed out. First, our study used only male rats. In prior studies using this discrimination procedure we have not observed large sex differences in responding (Strickland & McDannald, 2022; R. A. Walker et al., 2018). In our most recent study we constructed detailed, temporal ethograms for cue responding, finding females and males to be more similar than different (Chu et al., 2022). Still, it is possible that despite achieving similar behavioral discrimination different neural circuits may be recruited in females and males (Foilb et al., 2021).

Second, we used a human synapsin promoter that targeted all caudal substantia nigra cell types. So while co-expression of halorhodopsin and tyrosine hydroxylase was evident, it was not selective. This means we cannot conclusively link the effects of optogenetic inhibition to suppression of dopamine neuron firing. Indeed, the substantia nigra contains GABAergic projection neurons (Kirouac et al., 2004). We chose a pan-neuronal promoter for this study because our main question concerned the temporal specificity of caudal substantia nigra activity. Now that temporal specificity has been established, future studies can identify the cell types and projections underlying the function of the substantia nigra to suppress defensive responding.

Anatomical projections support the suggestion that either GABAergic or dopaminergic output neurons could contribute to nigral function in defensive settings. Nigra dopamine neurons project extensively to the forebrain and striatum where they are linked to reward learning (Waelti et al., 2001) and voluntary motor movements, among many other functions (Costa & Schoenbaum, 2022). In rat, the nigrostriatal pathway primarily targets the dorsal striatum. However, collaterals of striatal-projecting nigra neurons project to the central amygdala and ventral pallidum (Prensa & Parent, 2001). The central amygdala is viewed as a core brain region for defensive responding (Goosens & Maren, 2001; Koo et al., 2004; LeDoux et al., 1988; Li et al., 2013; McDannald, 2010; Moscarello & Penzo, 2022) and the ventral pallidum is emerging as a brain region controlling defensive and aversive behavior (Akmese et al., 2022; Correia et al., 2022; Farrell et al., 2021; Kaplan et al., 2020; Moaddab et al., 2021; Stephenson-Jones et al., 2020). Even more, substantia nigra GABAergic neurons project directly to the periaqueductal gray (Kirouac et al., 2004), a region long linked to defensive behavior (Fanselow, 1991; McNally, 2005; Rozeske et al., 2018; Shipley et al., 1991; Strickland & McDannald, 2022; P. Walker & Carrive, 2003; R. A. Walker et al., 2019; Wright & McDannald, 2019).

Here we have demonstrated that blocking caudal substantia nigra cue activity in a Pavlovian fear discrimination setting inflates defensive responding to uncertain threat and safety beyond control levels. These results capture a small segment of a core symptom of stress and anxiety disorders: excessive responding to non-threatening cues. Revealing the nigra projections and cell types that suppress defensive responding will broaden our understanding of defensive circuits and hasten the development of therapies to normalize exaggerated defensive responding to non-threatening cues.

## Acknowledgements

Research reported in this publication was supported by the National Institute of Mental Health of the National Institutes of Health under Award Number MH117791. The content is solely the responsibility of the authors and does not necessarily represent the official views of the National Institutes of Health. We thank Bret Judson and the Boston College Imaging Core for infrastructure and support. The authors report no competing interests.

